# Differential sensitivity of k-mer based identification tools using closely related database sequences affects relative abundance estimation

**DOI:** 10.64898/2026.10.15.759282

**Authors:** Annie J. Forster, Michael Payne, Rebecca J. Rockett, Cameron Webb, Tanya Golubchik

**Affiliations:** School of Medical Sciences and Sydney Infectious Diseases Institute, Faculty of Medicine and Health, The University of Sydney, Camperdown, New South Wales 2006, Australia; Centre for Infectious Diseases and Microbiology – Public Health, Westmead Hospital, Westmead New South Wales 2145, Australia; Department of Medical Entomology, NSW Health Pathology - Institute of Clinical Pathology and Medical Research, Westmead Hospital, Westmead, New South Wales 2145, Australia

## Abstract

Deep metagenomic sequencing is increasingly used for lineage profiling of complex biological mixtures, including applications such as wastewater virus surveillance and microbiome profiling. For profiling, sequence read sets are compared against a database of reference sequences to identify component genetic lineages, and estimate their relative abundance. However, there is currently a lack of clarity regarding the impact of two ubiquitous features of biological mixtures on compositional profiling: (1) population diversity, where reads contain genetic diversity not represented in the reference set, and (2) the presence of high sequence similarity between references, which causes ambiguity in read assignment.

Here we show, using both simulated and real data, that performance of two commonly used tools for lineage profiling, KMA and Kallisto, is differently affected by sample-reference distance and the presence of highly similar competing reference sequences. Increasing sample-reference distance reduced read assignment for both tools, but more strongly for Kallisto, which led to directional bias in abundance estimates when distance differed between mixture components. In contrast, when sample-reference distance was low, Kallisto more specifically discriminated between highly similar competing references, whereas KMA was less able to partition reads accurately between them. This specificity-sensitivity trade-off was reflected in real datasets: KMA produced more accurate abundance estimates for mosquito *COX1* mixtures, whereas Kallisto performed better for mixtures of closely related SARS-CoV-2 lineages, although neither tool fully resolved the most similar SARS-CoV-2 variants. Tool choice is therefore dependent on the degree of expected sample-reference distance and inter-reference distance within datasets.

## Introduction

With the advances of next generation sequencing, it is now possible to routinely produce large amounts of sequence data. This enables organism detection without the need for culture, and the ability to characterise complex genomic mixtures directly from their environment. This has many applications, including wastewater virus surveillance ^1^, microbiome profiling ^2^ and multi-pathogen screening ^3^.

For lineage composition analysis, large read sets are screened against a database of known reference sequences to identify the likely source lineage and estimate their relative abundance. However, accurate read assignment to each reference sequence poses challenges under two conditions. First, population diversity can mean that sequenced lineage reads do not match available reference sequences, preventing assignment to the expected reference. Second, reference databases may contain highly similar candidate sequences, causing ambiguity in assignment between competing references. These features are ubiquitous in real datasets and become particularly challenging when assessing under-sampled or highly diverse lineages.

A variety of tools can be used for reference-based read assignment. However, conventional mapping tools, such as BWA ^4^, do not explicitly handle multi-mapping to similar reference sequences. Pseudoalignment and k-mer based approaches, such as k-mer alignment (KMA) ^5^ and Kallisto ^6^, offer a computationally efficient alternative. These methods involve explicit decision-making to handle sequence similarity between reference sequences, such as favouring assignment to the reference with greater overall evidence (the ‘ConClave score’ implemented by KMA) or allowing reads to contribute to the support of multiple references (the Kallisto ‘equivalence class’). However, their performance under varying levels of inter-reference distance and sample-reference distance, and the implications of this for quantification of relative abundance in mixed samples, has not been completely characterised.

Here, we use simulations to isolate the effects of sample-reference distance and sequence similarity between references on read identification and relative abundance estimation performed by KMA and Kallisto. We then assess their performance in two contrasting use cases within real datasets of mosquito *COX1* gene and SARS-CoV-2 genome sequence data.

## Results

### Simulations isolate sample-reference distance and inter-reference distance

To isolate their effects on the performance of Kallisto and KMA, we designed simulations which separately modelled: (i) population diversity, by varying sequence divergence between the sample and reference (sample-reference distance), and (ii) the presence of highly similar competing reference sequences, by varying sequence divergence between reference pairs (inter-reference distance). Depending on the scenario, simulated reads from either a single source sequence or a mixture of reads from two source sequences were assigned to a reference set containing one or two sequences. Schematic trees depicting the relationships between sample component sequences and reference set sequences for each scenario are shown in **Figure S1**.

In each individual simulation, we generated a 10,000 base pair (bp) sequence of DNA (with even GC content) and randomly added single nucleotide substitutions (ranging from 1-2,000, at varying intervals). Reads simulated from these sequences (150bp paired end reads with an error rate of 1%, performed by wgsim) then underwent read assignment to the reference set using Kallisto or KMA, and the number of total reads assigned, and numbers assigned to individual references, were recorded. Ten independent sets of simulations were performed for each scenario.

In each of the following sections, we report the distance between sequences using mash distance which is based on the proportion k-mers which distinguish the two sequences (here computed using 31-mers; see **Methods**). A mash distance of zero means sequences share identical k-mer content, whereas values approaching 1 suggest little to no sharing of k-mers. Mash distance is strongly correlated with percentage identity difference but not in a one-to-one manner because it is dependent on the distribution of changes across overlapping kmers (**Figure S2**). We therefore use mash distance as the primary measure of sequence divergence, as it is likely to more closely reflect the decision making of Kallisto and KMA, both of which use k-mer based assignment.

### Sample-reference distance reduces single-lineage read assignment

First, we set out to test how increasing sample-reference distance affected the ability of each tool to assign reads from a single source to a single available reference. We found that increasing sample-reference distance reduced read assignment to the reference by both tools, but this was more pronounced for Kallisto (**Figure 1A**). As sample-reference mash distance increased to 0.075 (which in our simulations corresponds to up to 8% sequence mismatch across the 10,000bp sequence), Kallisto assigned 91.8% (95% confidence interval (CI) 90.7-92.9) of reads to the available reference, compared to KMA which assigned 97.8% (95% CI 97.5-98.2). This shows that Kallisto is more strongly affected by increasing sample-reference distance, resulting in progressive read loss as the sample grows more distant from the nearest reference.

**Figure 1.**
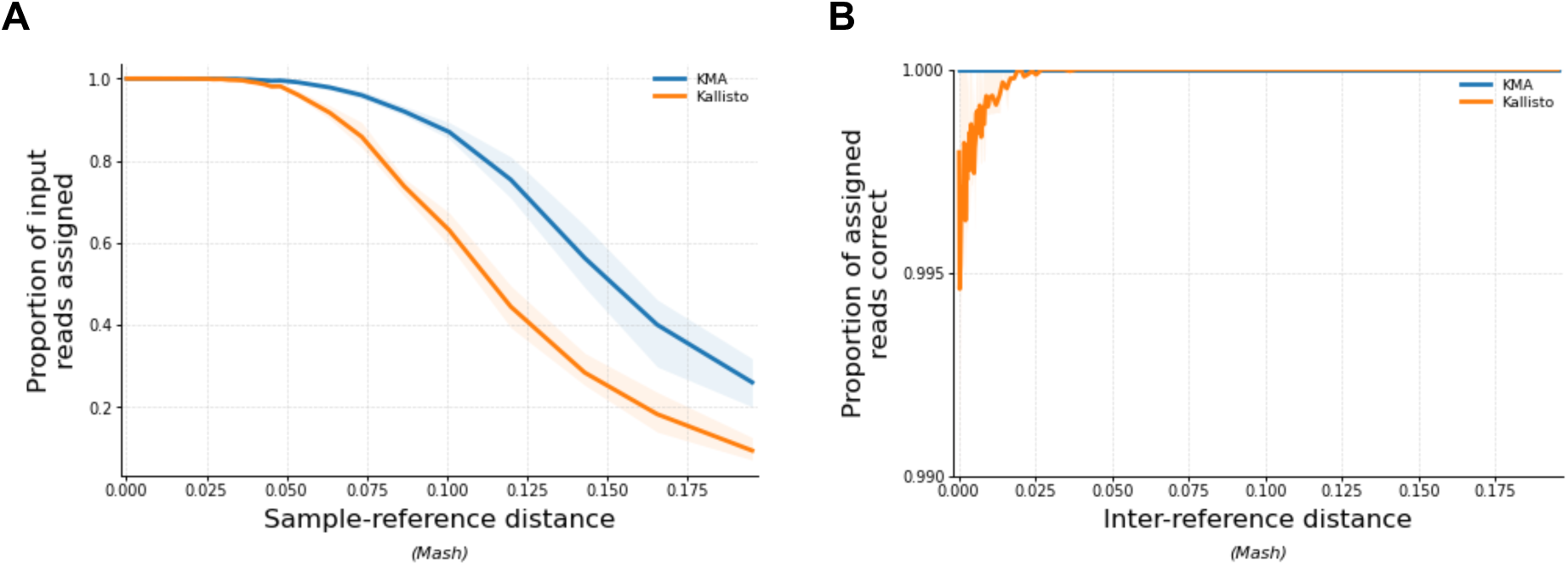
Effects of sample-reference distance and inter-reference distance on identification of reads from a single lineage. **A**, reads from a single source lineage were assigned to a reference set containing a single sequence while sample-reference distance was varied. **B**, reads from a single source lineage were competitively assigned to a reference set containing two sequences, one of which matched the reads perfectly, while inter-reference distance was varied. The x-axis shows the mash distance between the sample and the reference (A) or between the two available references.The y-axis shows the proportion of input reads which were assigned onto the single reference (A) and proportion of reads assigned to the perfectly matching available reference (B). All simulations used a 10,000bp sequence with 1–2,000 substitutions introduced at uneven intervals; reads were simulated and assigned against the relevant reference set. Each scenario was repeated 10 times, and lines show mean values across the repeats, computed across 75 quantile-based bins along the x-axis, where a minimum of 10 values were required to compute the average. Shading shows the interquartile range (25^th^-75^th^ percentile) of values within each bin. Results for Kallisto are shown in orange, and KMA in blue.

### Inter-reference distance affects single-lineage identification in the presence of sample-reference distance

Next, we tested how the presence of highly similar competing candidate references affects the ability of each tool to assign reads from a single source lineage to a perfectly matching reference. Both tools were able to assign the majority of input reads to the correct reference regardless of inter-reference distance, with a marginal reduction in accuracy for Kallisto at very low inter-reference distance; when mash distance between available references was 0.001-0.002 (equivalent to 10-20 mutations across the 10,000bp sequence), Kallisto assigned 99.7% of reads to the correct reference on average (**Figure 1B**).

We next tested whether inter-reference distance would show an effect when the tested sample was not perfectly represented in the reference (i.e. when sample-reference distance also varied). To model this, we used a tree structure where the sample belonged to a closely related sister clade of the nearest available reference (separated by a minimum mash distance of 0.01, i.e. 100 mutations; **Figure S1).** We then varied sample-reference distance and inter-reference distance simultaneously (shown on the x- and y-axes respectively in **Figure 2**). Under these conditions, decreasing inter-reference distance reduced the proportion of reads assigned to the correct reference for both tools (**Figure 2C**).

**Figure 2.**
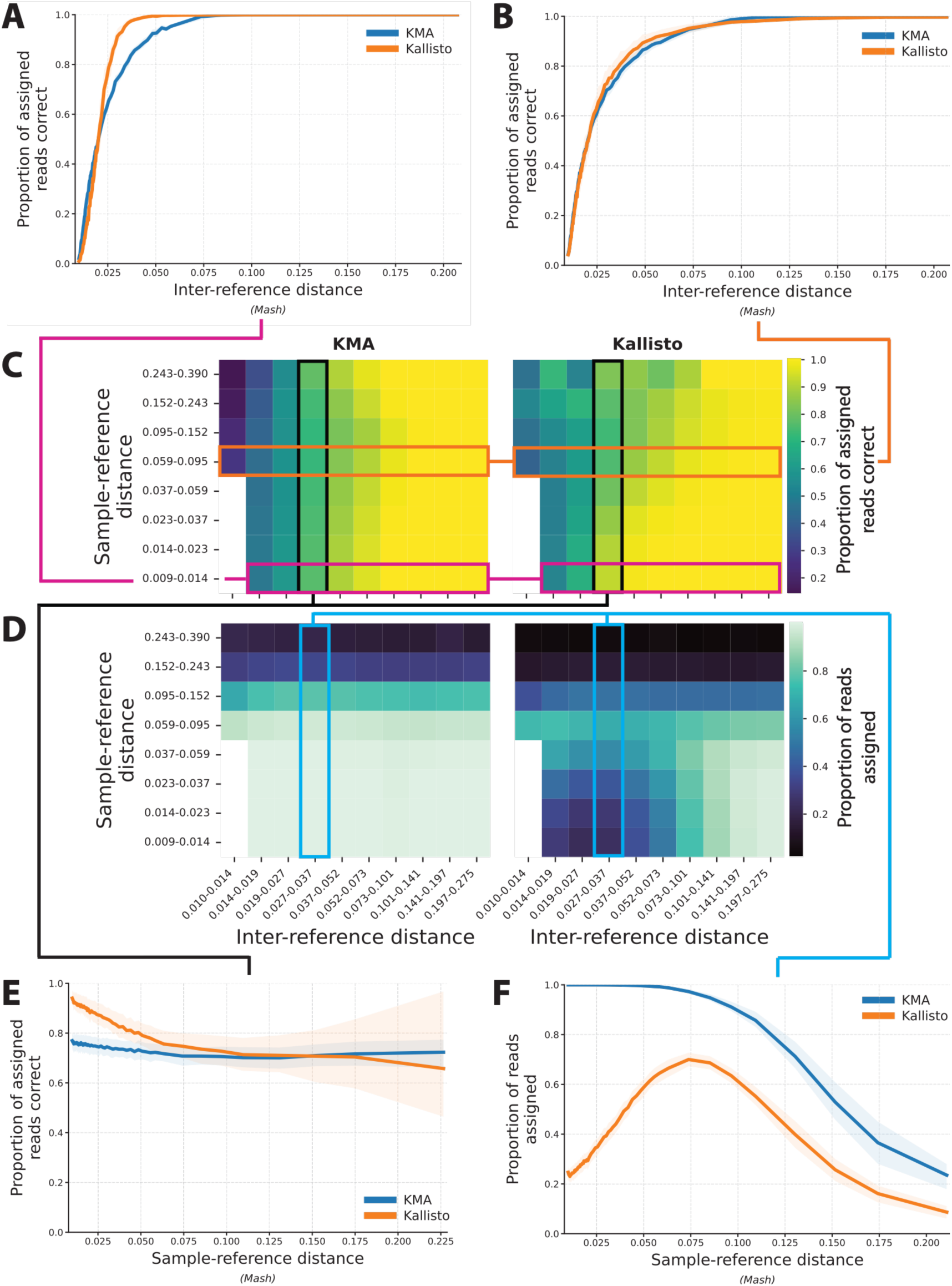
When no reference matches perfectly, sample-reference and inter-reference distance impact identification. In this simulation, reads simulated from a single source lineage were assigned to a reference set containing two sequences, while both inter-reference distance and sample-reference distance was varied. The sample belonged to a sister clade of its closest reference, separated by a minimum of 100 mutations. **C** and **D** show heatmaps across the full parameter space. Each row represents a range of sample-reference distance, and each column a range of inter-reference distance, as a mash distance. C shows the proportion of reads assigned to the correct reference of all assigned reads (specificity) where yellow indicates a high proportion and dark blue indicates a low proportion. D shows the proportion of input reads assigned (sensitivity) where light blue indicates a high proportion and dark blue indicates a low proportion. **A** and **B** show slices through panel C at low and moderate sample-reference distance respectively. When sample-reference distance was low (mash distance: 0.009-0.014), Kallisto more accurately discriminated reads between closely related references (A). When sample-reference distance was moderate (mash distance: 0.059-0.095), Kallisto and KMA performed similarly across reference distances (B). **E** and **F** show slices through panel C and D respectively at moderate inter-reference distance. When inter-reference distance was moderate (mash distance: 0.027-0.037), Kallisto more accurately discriminated reads when sample-reference distance was low (E). Kallisto initially excluded many reads at low sample-reference distance, followed by partial recovery when inter-reference distance increased (F). All simulations used a 10,000bp sequence with 1–2,000 substitutions introduced at uneven intervals; reads were simulated and assigned against the relevant reference set. Each scenario was repeated 10 times. For A, B, E and F, lines show mean values across the 10 simulations, computed across 75 quantile-based bins along the x-axis, where a minimum of 10 values were required to compute the average. Shading shows the interquartile range (25^th^-75^th^ percentile) of values within each bin. Results for Kallisto are shown in orange, and KMA in blue.

Tool performance depended on the degree of sample-reference distance. When sample-reference distance was low and inter-reference distance was low, Kallisto assigned a higher proportion of reads to the correct reference than KMA (e.g., when sample-reference mash distance < 0.02 and inter-reference mash distance was between 0.02-0.03, Kallisto assigned 71.4% (95% CI 71.1-71.8) of reads to the correct reference compared to KMA which assigned 62.5% (62.3-62.7) (**Figure 2A**). However, when sample-reference distance was high, KMA showed similar results to Kallisto (e.g., when sample-reference mash distance > 0.1 and inter-reference mash distance was between 0.02-0.03, Kallisto assigned 58.7% (95% CI 57.4-60.1) of reads to the correct reference versus 60.1% (95% CI 60.0-61.1) for KMA) (**Figure 2B**).

In this scenario, read exclusion by Kallisto was non-linear when inter-reference distance was low (**Figure 2D**). We attribute this to a process we have termed “k-mer collision”, where k-mers from the same read are attracted to different references and the read is discarded (drawn as a schematic in **Figure S3**). This leads to a reduction in read assignment by Kallisto at moderate sample-reference distance. However, as references become more distinct (i.e. inter-reference distance increases), some kmers become incompatible with any reference, allowing the read to be assigned again (e.g. **Figure 2F**). This helps explicate the observed patterns of Kallisto’s performance; by excluding ambiguous reads, Kallisto more specifically discriminates reads between closely related references when sample-reference distance is low, but its accuracy sharply declines as sample-reference distance increases (**Figure 2C**). In contrast, KMA excludes reads monotonically with increasing sample-reference distance, regardless of inter-reference distance (**Figure 2D**). Together, this indicates a trade-off between the specificity and sensitivity of these tools in the identification of reads.

### Low inter-reference distance impairs relative abundance estimation even when both sources match perfectly

Having established that both sample-reference distance and inter-reference distance impact identification of reads, we set out to assess their influence on relative abundance estimation. Here, relative abundance estimation was assessed using the log-ratio error between expected and estimated proportions between the two mixture components (shown in figures as fold-change).

We first tested how each tool was able to partition a set of mixed reads from two source lineages (in equal proportions) between two perfectly matching references. Under these conditions, low inter-reference distance impaired KMA’s ability to partition reads between the references. As inter-reference mash distance decreased to below 0.01 (corresponding to <1% sequence difference), the estimated ratio between components would diverge from 50:50 (e.g., at an inter-reference mash distance of 0.001, KMA absolute log-ratio error was 0.86, equivalent to a roughly 7-fold error in the inferred ratio between components, compared to an absolute log-ratio error of 0.01 for Kallisto) (**Figure 3**).

**Figure 3.**
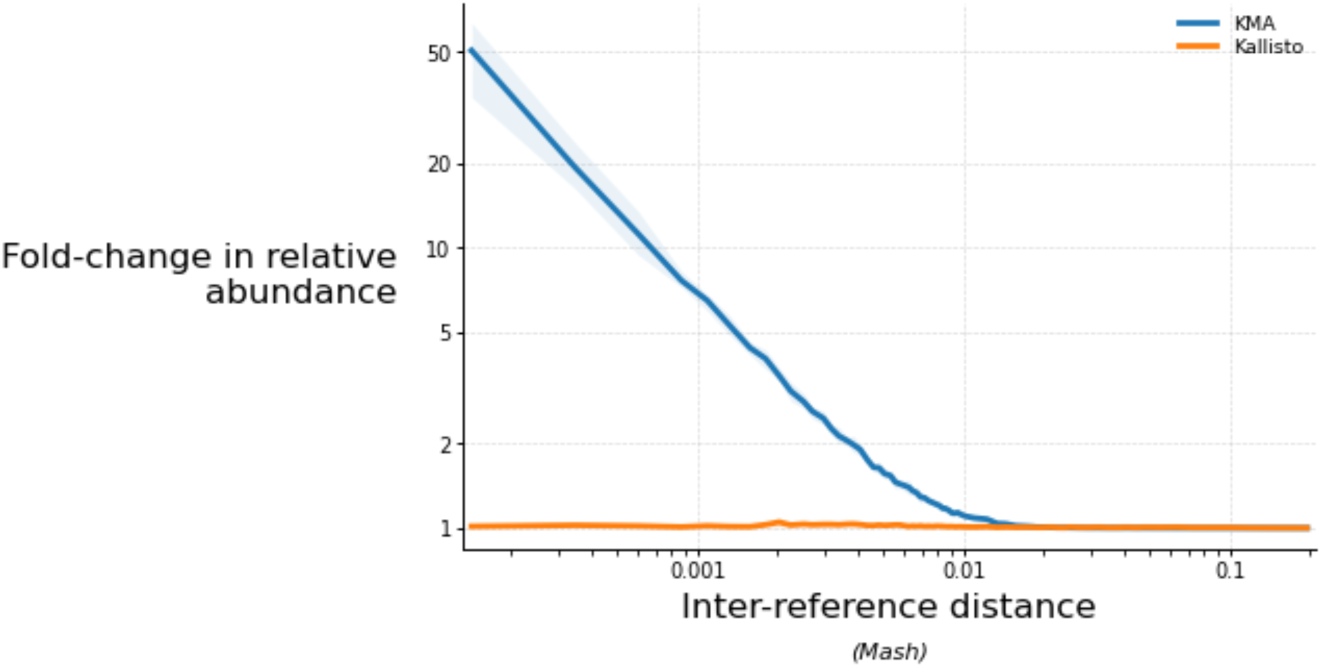
KMA is unable to partition reads from two lineages between very similar references. Reads from two source lineages were mixed in 50:50 proportions, and assigned to a reference set containing a pair of sequences which perfectly matched each source lineage, while inter-reference distance was varied. The x-axis shows the mash distances between the competing references. The y-axis shows the absolute fold error between the estimated and expected ratio of the two mixture components (computed as a log-ratio and displayed on a linear scale). All simulations used a 10,000bp sequence with 1–2,000 substitutions introduced at uneven intervals; reads were simulated and assigned against the relevant reference set. Each scenario was repeated 10 times, and lines show mean values, computed across 75 quantile-based bins along the x-axis, where a minimum of 10 values were required to compute the average. Shading shows the interquartile range (25^th^-75^th^ percentile) of values within each bin. Results for Kallisto are shown in orange, and KMA in blue.

### Asymmetric sample-reference distance biases relative abundance estimates

Next, we tested whether asymmetric sample-reference distance affected relative abundance estimation. In this set up, mixtures of reads from two source lineages (in equal proportions) were assigned between two references, one of which perfectly matched one component while sample-reference distance varied between the other sample-reference pair. This produced directional bias in both Kallisto and KMA, although this was more pronounced for Kallisto (**Figure 4).** For KMA, relative abundance estimation was accurate (absolute log-ratio error < 0.01) when sample-reference mash distance was within ∼0.07 (up to ∼750 substitutions in the 10,000bp sequence and ∼93% shared nucleotide identity), while Kallisto estimates were only unbiased when sample-reference mash distance was within ∼0.05 (roughly 550 separating substitutions, and 94.5-95% shared nucleotide identity).

**Figure 4.**
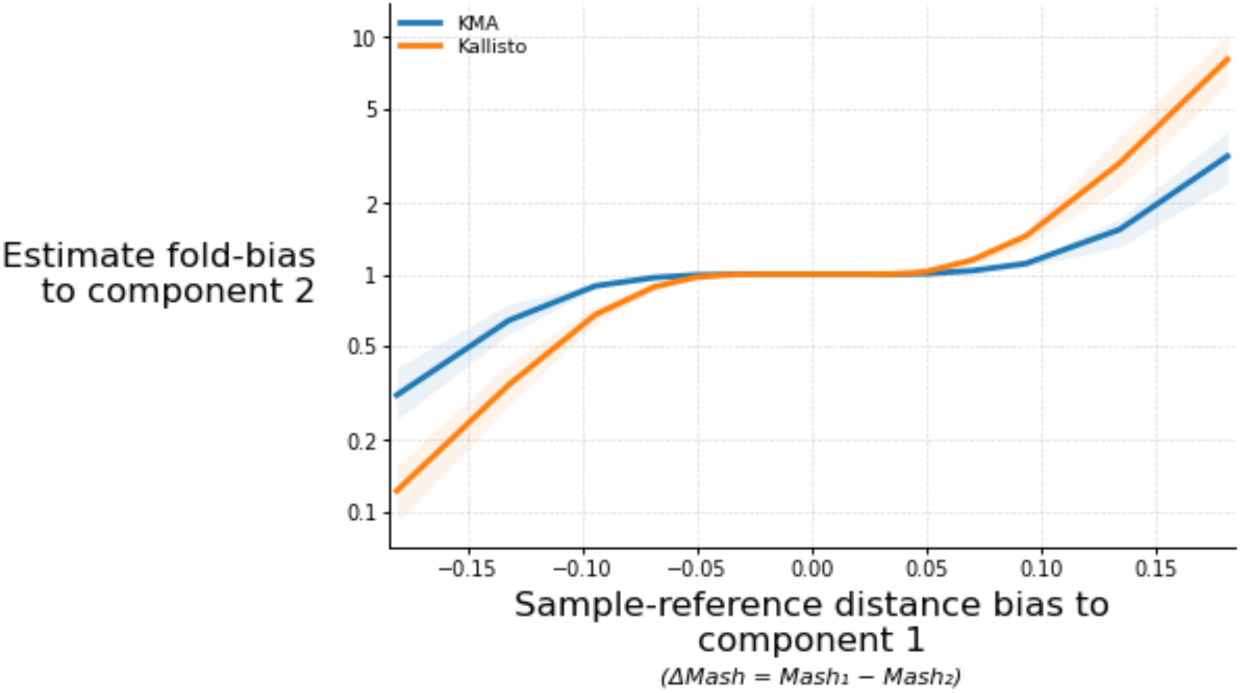
Asymmetric sample–reference distance biases abundance estimates toward the better-matching component. Reads from two source lineages were mixed in 50:50 proportions and assigned to a reference set containing a pair of sequences, where one sample-reference pair perfectly matched (i.e. sample-reference distance fixed at 0) and the other varied in sample-reference distance. The x-axis shows difference in sample-reference mash distance between the first and second reference; values <0 indicate that the second reference perfectly matched, while the first varied in sample-reference distance, while >0 indicates the converse. The y-axis shows the fold error between estimated and expected ratios of the two mixture components. All simulations used a 10,000bp sequence with 1–2,000 substitutions introduced at uneven intervals; reads were simulated and assigned against the relevant reference set. Each scenario was repeated 10 times, and lines show mean values, computed across 75 quantile-based bins along the x-axis, where a minimum of 10 values were required to compute the average. Shading shows the interquartile range (25^th^-75^th^ percentile) of values within each bin. Results for Kallisto are shown in orange, and KMA in blue.

### Extreme mixture proportions introduce an additional KMA bias

Finally, we set out to test whether the mixture proportions biases estimations made by these tools independently of sample-reference distance and inter-reference distance. In this test, reads from two sources were mixed in varying proportions and compared against a pair of perfectly matching references. We observed a marginal bias in KMA results to the minor component at extreme proportions (**Figure 5**). When the true abundance of a mixture component was more than 90%, KMA relative abundance estimates were biased by ∼1.05 fold on average toward the minor component (average absolute log-ratio error=0.022), whereas Kallisto showed little bias (average absolute log-ratio error=0.003).

**Figure 5.**
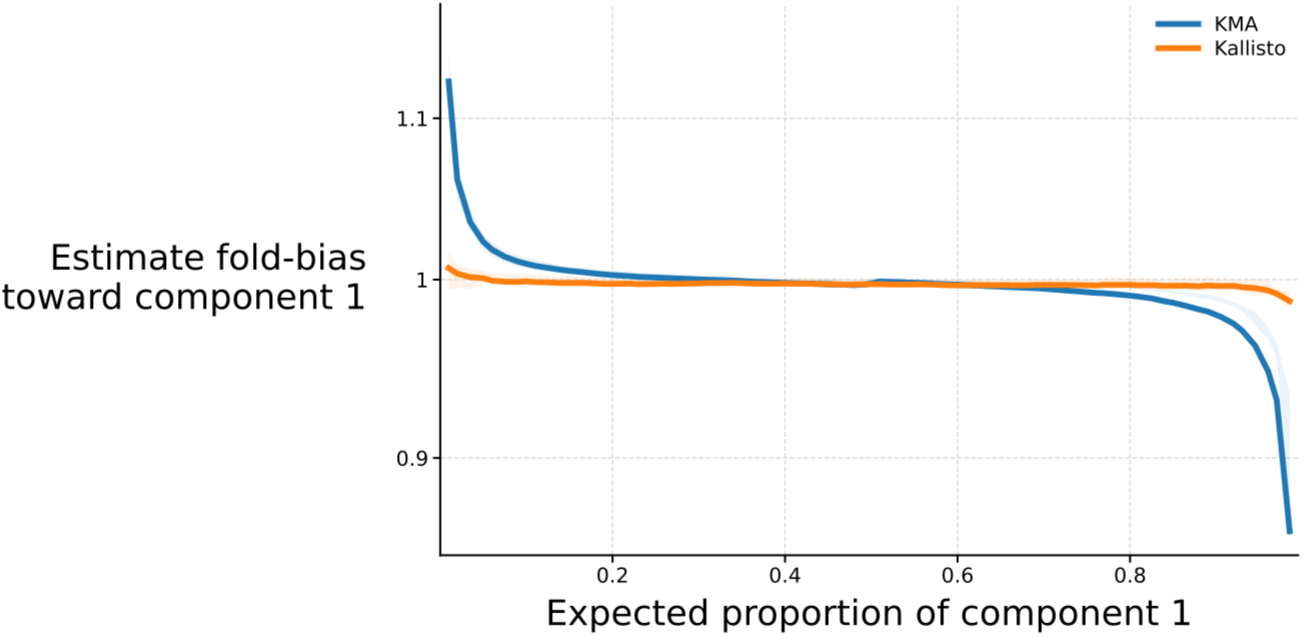
Extreme proportions bias KMA abundance estimates. Reads from two source lineages were mixed at varying proportions (1:99◊99:1, at 1% intervals) and assigned to a reference set containing a pair of sequences which perfectly matched the source lineage sequences (i.e. sample-reference distance was 0). Each sequence in the reference set was separated by 250 random substitutions. The x-axis shows the true mixture proportion. The y-axis shows the fold error between estimated and expected ratios of the two mixture components. All simulations used a 10,000bp sequence; reads were simulated and assigned against the relevant reference set. Each scenario was repeated 10 times, and lines show mean values, computed across 75 quantile-based bins along the x-axis, where a minimum of 10 values were required to compute the average. Shading shows the interquartile range (25^th^-75^th^ percentile) of values within each bin. Results for Kallisto are shown in orange, and KMA in blue.

### Real mosquito COX1 and SARS-CoV-2 data possess contrasting error characteristics

Real data varies by sample-reference distance and inter-reference distance. To assess how the behaviours identified by simulation impact real analysis, we selected two use-cases of contrasting characteristics: first, the deconvolution of mixtures of mosquitoes using the mitochondrial *COX1* gene, and second, the deconvolution of mixtures of SARS-CoV-2 lineages.

For each use-case, we compiled a reference set from a subset of available sequences. Mosquito *COX1* sequences were separated by large inter-reference distances, while SARS-CoV-2 lineages were genetically very similar, differing by as little as 9 polymorphisms (**Figure 6**). Conversely, sample-reference distance is expected to be greater for mosquito *COX1* sequences, where intraspecific diversity is likely to be incompletely represented in the reference set, compared to SARS-CoV-2 isolates which are typically similar to their assigned Pangolin lineage.

**Figure 6.**
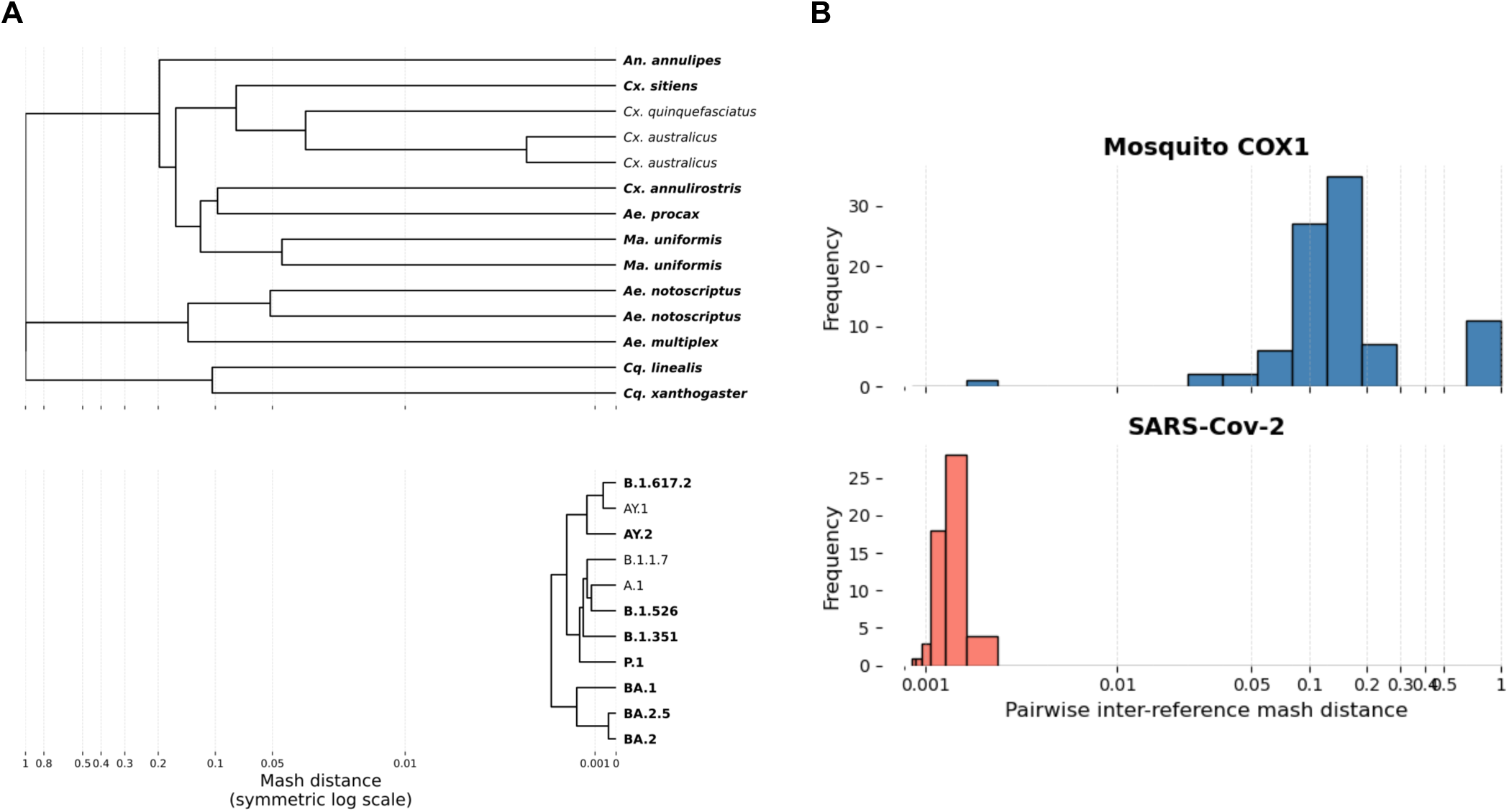
Relationships between reference sets of differing structure. Dendrograms (A) and histograms (B) showing the relationships between sequences in the mosquito *COX1* (top) and SARS-CoV-2 (bottom) reference sets based on Mash distance. The x axes are shown on a symmetric log scale). In A, species/virus lineages in bold denote those we possess sequence data for. Mash distance was computed between each pair of sequences (based on kmers of 31bp) and sequences were then clustered based on these distances using the complete method (i.e. based on the maximum distance between members of clusters).

### Real-data mixtures support simulation-derived tool trade-offs

We obtained *COX1* sequence data from 9 single-species pools of morphologically identified mosquitoes and publicly available amplicon data from 8 SARS-CoV-2 samples (**Table S1**). We then generated a dataset of pairwise mixtures of sample reads *in silico* (in 10:90, 25:75, 50:50, 75:25 and 90:10 proportions) and estimated composition using Kallisto and KMA.

First, we assessed how well each tool identified the mixture components. Both tools often struggled to correctly identify the SARS-CoV-2 components, consistent with the extremely low inter-reference distances in the reference set (**Figure 7A).** On average, KMA assigned 20.8% (95% CI 15.3-26.4) of reads to incorrect SARS-CoV-2 references, while Kallisto assigned 29.1% (95% 25.5-32.6). For the mosquito *COX1* dataset, KMA misidentified reads slightly more often than Kallisto, assigning 3.8% (95% 3.1-4.5) compared to 1.8% (95% 1.4-2.2) (**Figure 7A**), indicating lower specificity of this tool.

**Figure 7.**
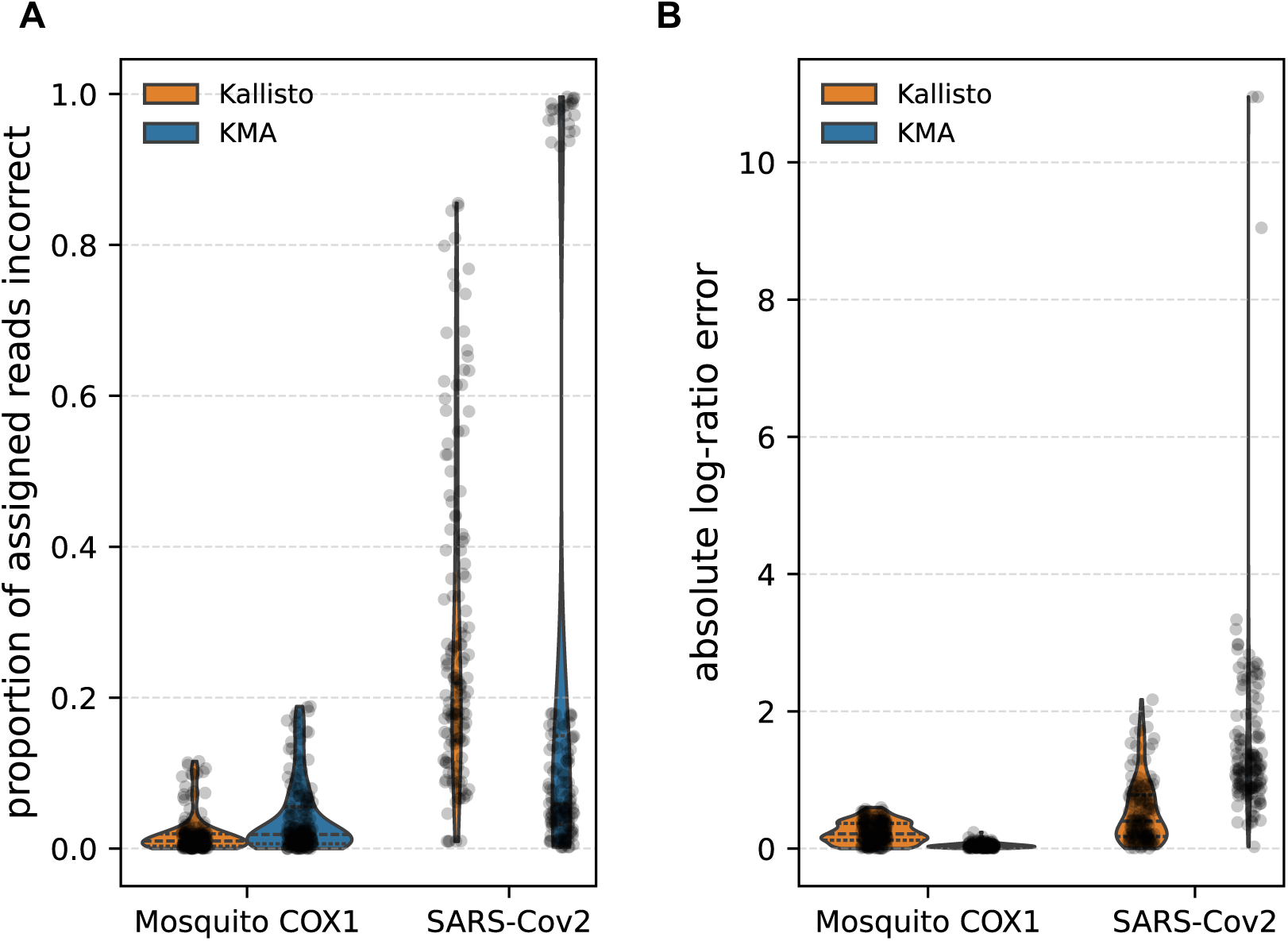
Identification and relative abundance error across mixtures of real data. A shows the proportion of all assigned reads that were assigned to an incorrect reference (y-axis) for the mosquito *COX1* and SARS-CoV-2 datasets (x-axis). B shows the absolute log-ratio error between the expected and estimated proportions (y-axis) between the datasets. Results are depicted as violin plots which are lined to show the median value (dashed) and inter-quartile ranges (dots). The actual computed values have also been displayed on the violin plots as dark gray points. Results for Kallisto are shown in orange, and KMA in blue.

However, KMA gave more accurate relative abundance estimates between the true components for the mosquito *COX1* mixtures (KMA average absolute log-ratio error=0.05 (95% CI 0.040-0.050), Kallisto average absolute log-ratio error=0.23 (95% CI 0.206-0.253); **Figure 7B**). This is consistent with Kallisto’s greater sensitivity to sample-reference distance (**Figure 2B**). Kallisto estimated the relative proportions of the two SARS-CoV-2 components more accurately (average absolute log-ratio error=0.64 (0.47-0.80) compared to 2.94 (95% CI 2.36-3.51) for KMA)), matching the earlier findings that suggest Kallisto is better able to partition reads among similar references.

However, neither tool resolved these SARS-CoV-2 mixtures well in absolute terms because they each assigned a large proportion of reads to incorrect references. This suggests that, for such datasets, reference clustering at a higher taxonomic or nucleotide identity sharing level may be necessary to obtain more reliable abundance estimates.

## Conclusions

Reference-based read assignment is widely used in genomics. Using this to estimate relative abundance is a natural extension to this and is particularly desirable within the fields of meta-genomic and meta-transcriptomic sequencing. However, accurate abundance estimation depends on how well the sampled sequences are represented in the reference set and on how distinguishable the competing references are from one another. Here, we show that these two properties impact results produced by the two tools, Kallisto and KMA, in different ways and to different degrees.

Kallisto appears to be more sensitive to sample-reference distance, tending to discard reads at higher rate than KMA (**Figure 1A**). Simulations suggest that when sample-reference distance was low, Kallisto was more able to discriminate reads between more similar references than KMA (**Figure 2**). KMA on the other hand, was more robust to sample-reference distance; when sample-reference distance was present, KMA performed as well as Kallisto across inter-reference distance ranges.

This is likely due to the algorithmic basis of each tool. Kallisto locates exact kmer matches between read and reference sequences. If reads are compatible with multiple references (known as an equivalence class) they can contribute to the evidence for either reference. However, when separate kmers from a single read align to different references, which can occur due to a combination of high sample-reference distance and low inter-reference distance, then reads are discarded. We refer to this as kmer collision (**Figure S3**).

KMA initially seeds alignments based on kmers and builds out from this to generate a score which is compared to other possible alignments to determine read assignment. Every read, given it has a kmer match to a reference, can therefore be assigned leading to greater read use by this tool. However, small differences between scores, such as due to single nucleotide polymorphisms, may not sufficiently discriminate between very similar references, leading to reduced specificity by KMA.

Taken together, this displays a sensitivity-specificity trade-off; Kallisto favours specificity by discarding ambiguous reads, while KMA is more sensitive to the detection of more distantly matching samples.

This has knock on effects for data analysis and suggests the best approach to relative abundance estimation depends on the likely sources of error in the dataset. Where targeted taxa are under-sampled, sample-reference distance may vary between sample-reference pairs leading to biases in estimates (**Figure 4**). In an example use-case, we imagined estimating the relative abundances of mosquitoes in multispecies pools by mixing *COX1* gene reads from wild-caught mosquitoes *in silico*. References were dissimilar (i.e. high inter-reference distance), and sample-reference distance is likely to be high, and as such KMA produced more accurate abundance estimates between pairs (**Figure 7**). In a contrasting scenario, we imagined resolving very closely related SARS-CoV-2 lineages also from *in silico* mixtures of amplicon data. Sample-reference distance was likely to be minimal, favouring Kallisto, although these mixtures were not well-resolved with either tool, suggesting there is a lower bound to inter-reference distance that is currently possible to resolve with these tools.

In conclusion, tool choice for relative abundance estimation depends on expected dataset characteristics. If samples are likely to be diverse compared to available references and references are sufficiently distinct (e.g. mosquito *COX1*), KMA may be more suitable. If samples closely match highly similar references (e.g. SARS-CoV-2 lineages), Kallisto may be more accurate. Sample-reference distance may be challenging to control in real analyses, but inter-reference distance may be fine-tuned by clustering similar references to within a discriminable limit.

## Methods

### Simulated data

#### Simulation design overview

We set out to design a simulation framework which could independently vary (i) inter-reference distance and (ii) sample-reference distance, in order to isolate their individual effects on results produced by the read assignment tools Kallisto^6^ and KMA^5^. In total, we tested five different scenarios as well as a sixth scenario that tested biasing due to mixture proportions. These were:

i. How varying sample-reference distance affects the assignment of a single source of reads to a single reference (**Figure 1A**)
ii. How varying inter-reference distance affects the assignment of a single source of reads between two references (**Figure 1B**)
iii. How varying sample-reference distance and inter-reference distance affects the assignment of a single source of reads between two references (**Figure 2**)
iv. How varying inter-reference distance affects the partition of an equal mixture of reads between two references (**Figure 3)**
v. How asymmetric sample-reference distance in an equal mixture of reads affects relative abundance estimations between two references (**Figure 4**)
vi. How mixture proportion affects the partition of a mixture of reads between two references (**Figure 5**)

Scenarios i-iii were designed to test the impacts of inter-reference distance and sample-reference distance on read identification, whereas iv-vi were targeted to assess relative abundance estimation. The first three scenarios therefore involve a single source of reads, whereas the latter three involve two read sources. Overall, we performed simulations for each scenario ten times, and displayed results show an average across the ten replicates.

#### Sequence generation and mutation scheme

For each simulation, we simulated a 10,000 bp sequence (using the Python package Numpy (v2.3.4)) with even GC content, and randomly inserted 1–2,000 randomly placed single nucleotide mutations. The numbers of inserted mutations were such that there was an interval of 1 mutation between 1-100, 5 mutations between 100-250, 10 mutations between 250-500, 50 mutations between 500-1000 and 100 mutations between 1000-2000 (totaling 175 sequences in addition to the original). Varying intervals were used in this way to speed up computation, and because during initial exploration we noted that read identification and relative abundance estimation changed most rapidly at low numbers of mutations and then plateaued at higher levels of divergence. Therefore, more granular intervals were used at low mutation numbers where changes in behaviour were greatest. Mutations were inserted with replacement, and it is therefore possible that mutations were reverted, however we did not deem this problematic as this would be reflected in the Mash distance used for assessment of differences between sequences (see below).

#### Read simulation

From each sequence simulated as described above, 1,000 read pairs of 150bp length were simulated using wgsim^7^ (v0.3.1-r13). We included an error rate of 1%. These were stored and then either run through the assignment tools KMA and Kallisto against a particular reference or mixed into a new file as described below.

### Real datasets

To test Kallisto and KMA in a more real data analysis scenario, we compared two different use-cases. In the first, we assessed using Kallisto and KMA for identifying mosquito species and estimating their proportional abundances in sequence data generated from a pooled collection, reflecting a potential ecological surveillance scenario utilising the mitochondrial *COX1* gene. In the second, we assessed using the tools to identify and measure the relative abundance of closely related SARS-CoV-2 lineages, simulating wastewater surveillance.

In brief, we located identified reference *COX1* gene sequences and SARS-CoV-2 whole genomes, and compiled these into two reference sets. We then took short-read sequence data generated for the mosquito *COX1* (from single species pools), and amplicon short read data for different SARS-CoV-2 lineages. We then mixed sequence reads *in silico* and compared estimated relative abundance estimation by each tool compared to the true mixture. Further details of these steps is given below.

#### Reference set construction

To test sequence data for the mosquito *COX1* locus, we first constructed a database of references which included the nine species we acquired short read sequence data for (listed in **Table S1**). As part of a separate study, we had previously compiled a larger reference dataset aiming to cover species present within Australia. Because this covered more than 70 species, we reduced this for the purposes of the tests conducted here to include the nine test species, and two additional related species common in Australia (*Culex australicus* and *Cx. quinquefasciatus),* totalling 11 species represented in the reference.

To find these sequences, NCBI was manually searched for a mitochondrial genome or *COX1* gene sequence. Partial *COX1* sequences which have been generated from sanger sequencing are commonly available. These are often of <700bp in length (the mosquito *COX1* gene is roughly ∼1,500bp in length). However, through initial testing of Kallisto and KMA, we found that the presence of portions of *COX1* present in some samples and missing in others lead to the assignment of reads to where the region was present, leading to inaccurate calls. We tackled this in two ways. First, we set a minimum length threshold of 1,000bp for *COX1* sequences during our manual NCBI search, and second, we opted to trim sequences to homologous regions of the gene. To do so, we generated a multiple sequence alignment (MSA) using MAFFT (v7.526)^8^ and using custom code, we scanned across the MSA and trimmed regions from either end where consecutive gaps occurred across the set. A small gap allowance of 3bp was permitted within each sequence to allow for INDELS within the sequence. This left a 930bp core region shared across available *COX1* sequences in the database. If a species had multiple near complete sequences available at the time of searching, we retained both (*Mansonia uniformis* and *Cx. australicus*) we retained both sequences. 14 individual sequences were therefore included in the reference.

Although many lineages of SARS-CoV-2 have been identified since its origin, we also focussed on a limited set of 11 lineages here. These were chosen prior to short-read data acquisition and were initially selected based on a benchmarking dataset released by Aßmann *et al.* (2024)^9^, although we do not make use of this data here. Instead, whole genomes for these lineages were searched for on NCBI Virus by filtering for complete genomes with no missing nucleotides and downloaded in FASTA format from NCBI. For common lineages, many hundreds of complete genomes are available, so we chose to include one per lineage. Lineages and accessions are listed in **Table S1.**

#### Mosquito *COX1* sequence data generation

Mosquito specimens were sourced from field collections at Sydney Olympic Park. Mosquitoes were collected from carbon dioxide baited light traps and identified to species using taxonomic and pictorial guides. Identification was performed by experts within New South Wales Health.

Mosquito sequence data was generated via a targeted metagenomics sequencing approach which included probes to enrich the mosquito *COX1* gene. Specimens were combined into nine same species pools (species listed in **Table S1**). DNA was extracted from pools, and library preparation was performed using the EF Library preparation kit (version 1, Twist Biosciences). Following standard library preparation, capture was performed on libraries, and these were sequenced on the Illumina NextSeq machine at 150bp paired end using 300 cycles.

#### SARS-CoV-2 amplicon data collection

In order to broadly match the produced mosquito *COX1* sequence data, we sought out paired-end sequence data for the SARS-CoV-2 virus. The vast majority of available data for this virus has been produced by an amplicon-based protocol which tends to produce near complete coverage of the genome ^10^. We sought out data for each tested lineages manually by filtering by the Pangolin lineage and including an accession column on the NCBI virus website, and inputting returned accessions into ENA. Once ensuring that two files were available for download, representing paired end data, the raw data was downloaded.

#### Sequence data pre-processing

As outlined above, the sequence data produced from the single species mosquito pools underwent a targeted meta-genomics sequencing approach which targeted the *COX1* gene and arbovirus sequences. It is therefore possible that sequence data represents other taxa than the mosquito alone. Before read assignment we therefore decided to filter data to approximately represent mosquito *COX1* sequences alone. We also treated the SARS-CoV-2 amplicon data in a similar way.

These steps were as follows. First, we set out to generate a sequence which represents diversity across the mosquito *COX1* and SARS-CoV-2 sequences. For the mosquito *COX1* we used the reference set described above, whereas for SARS-CoV-2, the RefSeq sequence was used (Wuhan-Hu-1; NC_045512.2). For each of these reference sets, sequences were clustered using CD-HIT^11^ to within 90% sequence identity. A consensus was then generated from each cluster using viralconsensus^12^ default parameters to determine the consensus nucleotide at each position. For the mosquito *COX1* reference set this produced three sequences which roughly align to the *Anopheles*, *Aedes* and *Culex* genera, while SARS-CoV-2 produced one sequence.

Sequence reads were then aligned to these consensus sequences. For SARS-CoV-2, minimap2 (v2.30-r1287) was used, and for mosquito *COX1* BWA was used as implemented by the Castanet tool ^13^. All mapped reads were then extracted using samtools^7^ (v1.22.1) (secondary and supplementary alignments were not removed) and converted to FASTQ format using bedtools^14^ (v2.31.1). These reads were then input into read assignment using Kallisto and KMA.

#### *In silico* mixture generation

In order to test relative abundance estimation for the simulation analysis (scenarios iv-vi described above), and in the assessment of mosquito *COX1* and SARS-CoV-2 relative abundance estimation, simulated and real sequence reads were mixed *in silico*.

For the simulated sequence data, producing a mixture of equal proportions (i.e. simulation scenarios iv and v) was performed using the bash command “cat” to concatenate reads from two sources into a single file. As such all 1,000 simulated read pairs were combined into the same FASTQ file (totalling 2,000 read pairs).

For simulation scenario vi, depicted in **Figure 5**, we wanted to ensure sequence coverage would not impact results. In this scenario we therefore computed the minimum number of reads which would roughly correspond to 5-fold coverage of the sequence. We simulated extra reads and mixed reads to proportions varying from 1% to 99% (at 1% intervals) where the number of reads in each proportion could not be less than the minimum read value. In this case, reads were mixed using the bash command “head” with the flag “-n” to select the desired number of reads (four lines per read are present in a FASTQ file).

For the mosquito *COX1* and SARS-CoV-2 analysis, the program seqtk^15^ (v1.5-r133) and subcommand ‘sample’ was used to randomly sample sequence read pairs and place them into a mixed file. The total numbers of reads to sample from each component was determined using the minimum total read count in either of the source files to ensure desired proportions would be satisfied. In these analyses, reads with mixed in the ratios 10:90, 25:75, 50:50, 75:25 and 90:10 for every possible pair in the set.

### Tool execution

In this study, we focus on the tools Kallisto and KMA for read assignment to references. Both of these first build an index file for the input reference set. For our simulation analysis, many references were generated which either contained one (e.g. simulation scenario i) or two sequences (e.g. simulation scenarios ii-vi). These were generated by first combining sequences into a single FASTA file using the bash command “cat”, and using the tools as described below.

In the mosquito *COX1* analysis, some mosquito species had multiple representatives in the reference set. In this case, we would sum read counts across the species for Kallisto and KMA to get a total for that species.

#### Kallisto

Kallisto (v0.51.0) index generation (the “index” subcommand) and read assignment (the “quant” subcommand) were run with default command line arguments. Read assignment in this study was performed without the use of bootstrapping. Read counts on individual references were then determined by the “est_read_counts” column in the resulting abundance.tsv file.

#### KMA

KMA (v1.6.8) index generation (the “index” subcommand) was run using default command line arguments. Read assignment was run in paired end mode (“-ipe”) using the additional command line arguments “-ef -ex_mode -vcf 2 -1t1”. The numbers of reads assigned to each reference was determined by dividing the “readCount” column in the produced “.mapstat” file by 2 (in order to report read pair numbers, as all data used in this manuscript are paired end).

### Distance and performance metrics

#### Mash distance

Because Kallisto and KMA are kmer based tools, we chose to report results in terms of the Mash distance between sequences. Mash is derived from the jaccard distance between two sequences which is the proportion of shared kmers out of all kmers between sequences. When mash is zero, this indicates that sequences are identical in terms of their k-mer content, and as values approach 1 this indicates little to no shared kmer content. Mash distance is strongly negatively correlated with percentage identity sharing between sequences (**Figure S2**), but more effectively accounts for dispersal of mutations across the sequence.

#### Measuring identification accuracy

Identification was measured by determining the proportion of input reads which were assigned to the correct reference, where correct means the closest relative to the source of the reads. For simulation scenarios testing sample-reference distance this will usually not be an identical match.

For the real data comparisons, the correct reference was considered that which matches the morphological (mosquito *COX1*) or Pangolin lineage (SARS-CoV-2) identification given to each sequence. Complete details are given in the respective figure legend of each presented analysis.

#### Measuring relative abundance performance

Error in relative abundance estimates were measured by comparing the expected and computed mixture proportions. Let a mixture of sequence reads contain reads from taxa *A* and *B*, and *q_A_* and *q_B_* represent the true mixture proportions of components A and B. *q̂_A_* and *q̂_B_* represent the estimation made by KMA or Kallisto. The relative log-ratio error between these is therefore:

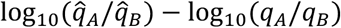

The value is a signed measure of bias in the estimated proportions where negative values indicate the estimation has been biased towards B and positive values indicate towards A. When converted to linear space, this equates to the fold error in the ratio of component A to component B relative to the true ratio, where values <1 indicate bias towards B and values >1 indicate bias towards A. For analyses concerned with magnitude rather than direction of estimation error, we report the absolute log-ratio error. To prevent zero values from producing log errors, a small value (10^-10^) was added to both the true and estimated ratio.

Mean values are reported with 95% confidence intervals calculated as the mean ± 1.96 standard errors.

## Data and code availability

Code associated with this manuscript is available on GitHub (https://github.com/AnnieJForster/KMA-kallisto-comparison-paper) with data files available for download from [Zenodo](https://zenodo.org/records/23328846) (10.5281/zenodo.23328846). Mosquito sequence data will be made available from the European Nucleotide Archive (ENA) with project ID PRJEB128300.

## Supplementary

**Figure S1.**
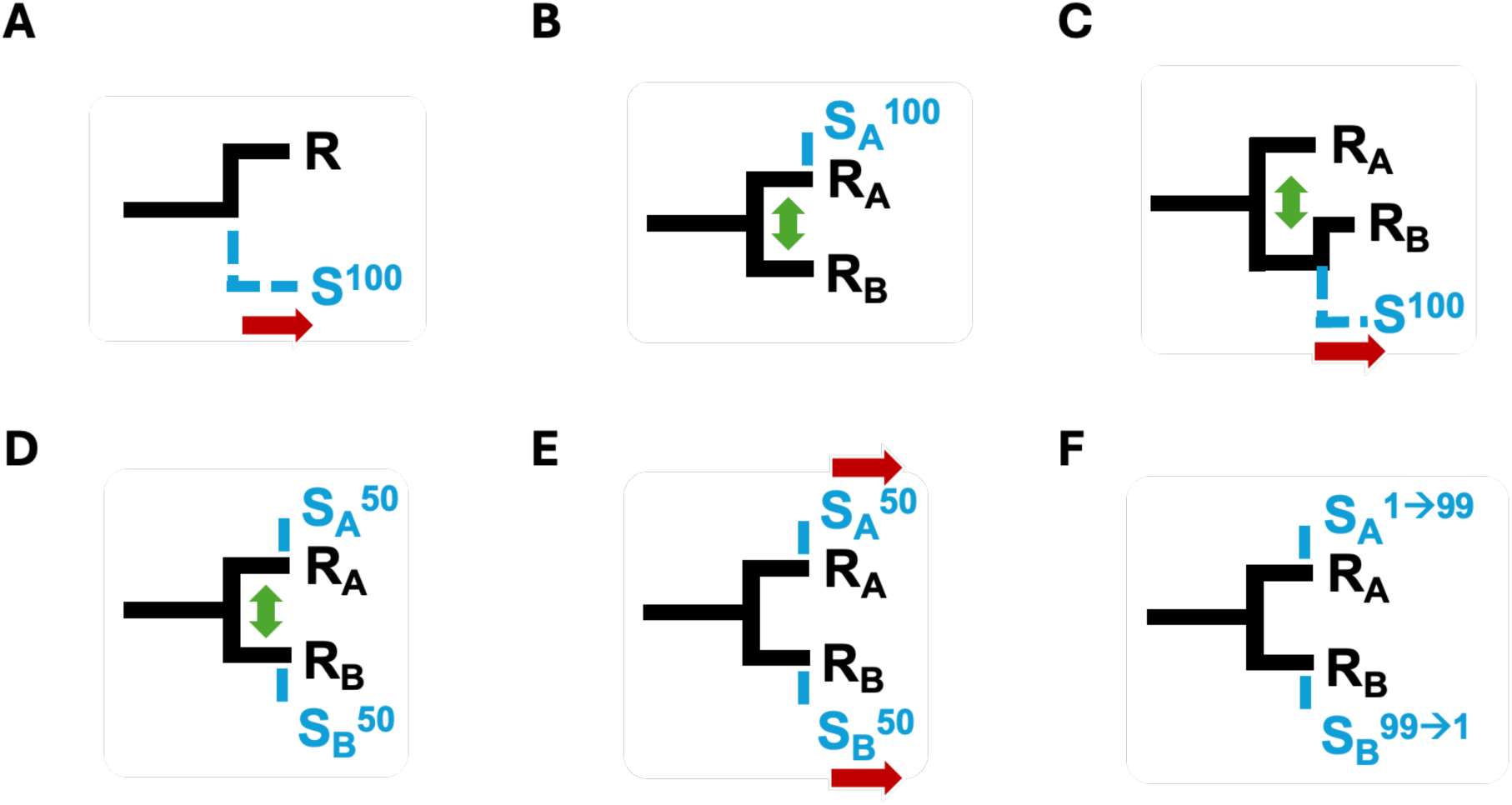
Schematics of the six tested simulation scenarios presented in this paper to test the performances of Kallisto and KMA. Panels A-F correspond to scenarios i-vi described in Methods. In all panels, black solid lines and text indicate the references used in read assignment, while blue dashed lines and test indicate the samples tested on those references. References are named “R” while samples are named “S”. If multiple sequences were included in the reference, the subscript gives a label to that reference and sample (A or B) showing they are the closet related pair. If a sample contained a mixture of reads from two sources, the percentage proportion of reads belonging to that sample is shown as superscript. Large red arrows indicate that in the simulated scenario, sample-reference distance was varied, while double-headed green arrows indicate inter-reference distance was varied. In panel E, sample-reference distance between R_A_ and S_A_ or sample-reference distance between R_B_ and S_B_ was varied at one time. In panel F, rather than sample-reference distance or inter-reference distance, mixture proportion was varied indicated by arrows within the superscript.

**Figure S2.**
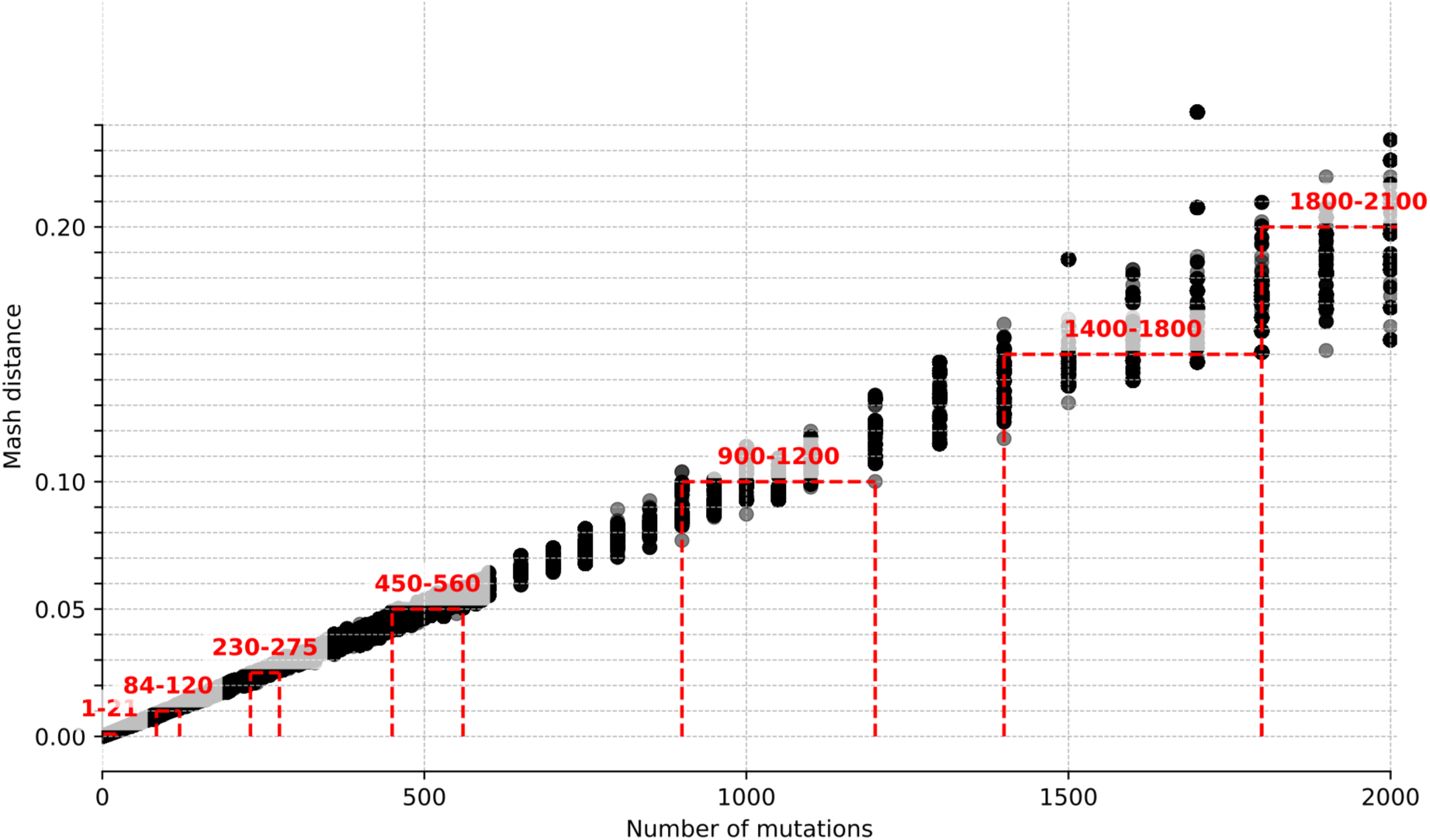
Relationship between Mash distance and number of mutations inserted a 10,000bp sequence. The x-axis shows the number of mutations inserted into a sequence and the y-axis shows the Mash distance between the original and mutated sequence. These show values from across the simulation scenarios displayed in this paper. To help interpret values presented in the paper, ranges of numbers of inserted mutations corresponding to a Mash distance of 0.001, 0.01, 0.025, 0.05, 0.1, 0.15 and 0.2 are shown in red, and the range of mutation numbers this corresponds to annotated.

**Figure S3.**
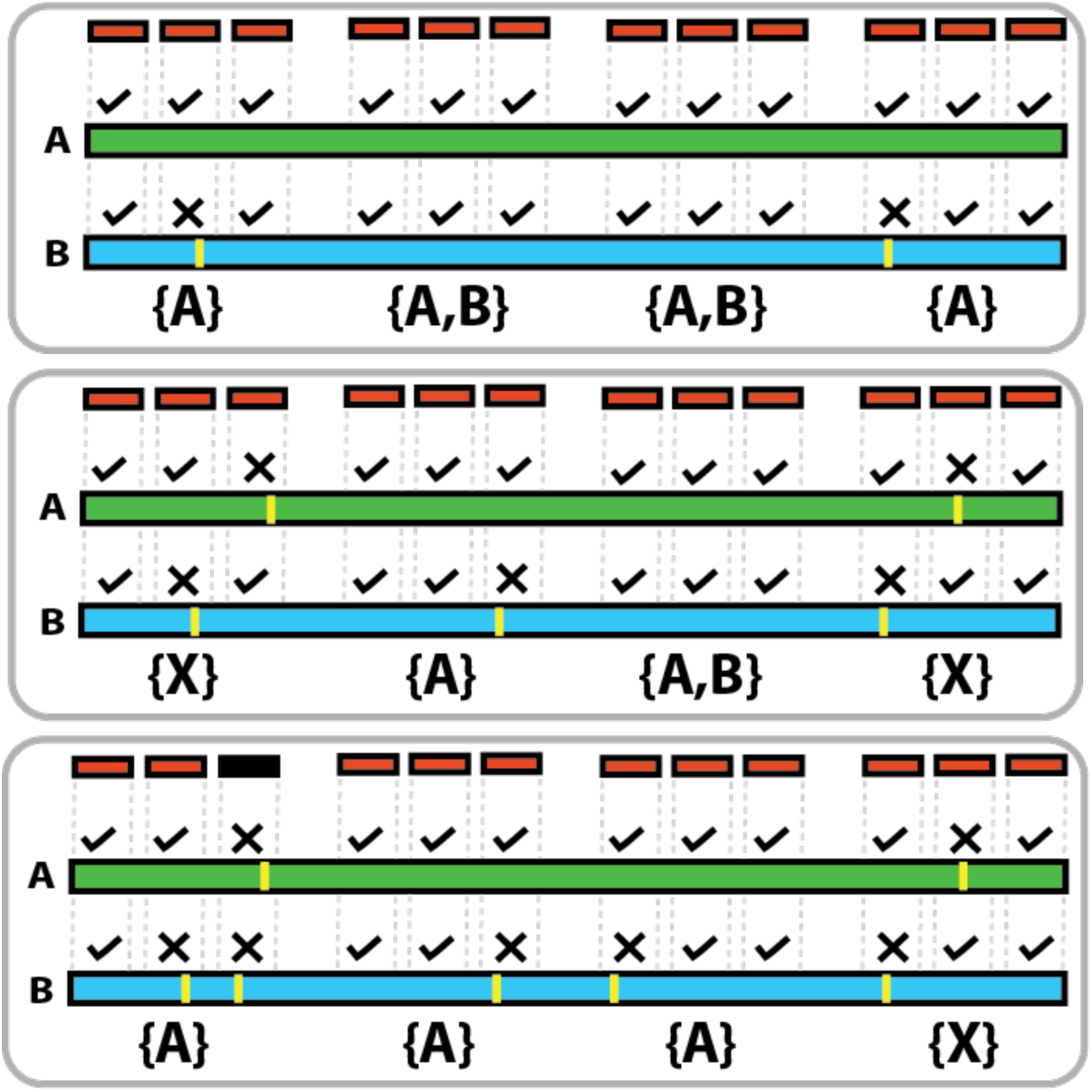
Diagram of how k-mer collision impacts read assignment by Kallisto. Each read contains three k-mers (red). A read can be assigned only if all compatible k-mers support the same reference. Assignable kmers to an individual reference are marked with a tick, while those that can’t are marked with a cross. If a read is compatible with multiple references, it forms an equivalence class (e.g, “[A,B]”). Top; when one reference matches the sample, reads are assigned correctly. Middle; when neither reference matches well, k-mers from the same read may support different references, so the read is discarded (“[X]”). Bottom; when the references are more distinct, some sample k-mers are incompatible with any reference (black), allowing the read to be assigned.

**Table S1.** Table of accessions for reference sequences and sequencing data used in this manuscript. Where data type is ‘Genome’, these sequences contributed to the reference, while amplicon and single species pool data were used as read sources to test assignment.

| *Taxa/virus* | *Species/lineage* | *Data type* | *Accession* |
| --- | --- | --- | --- |
| SARS-CoV-2 | A.1 | Genome | PQ540440.1 |
| SARS-CoV-2 | B.1.1.7 | Genome | BS016086.1 |
| SARS-CoV-2 | B.1.351 | Genome | PV241309.1 |
| SARS-CoV-2 | P.1 | Genome | PX578267.1 |
| SARS-CoV-2 | B.1.526 | Genome | PV001695.1 |
| SARS-CoV-2 | B.1.617.2 | Genome | PX952986.1 |
| SARS-CoV-2 | AY.1 | Genome | OR363636.1 |
| SARS-CoV-2 | AY.2 | Genome | PQ982016.1 |
| SARS-CoV-2 | BA.1 | Genome | PX503961.1 |
| SARS-CoV-2 | BA.2 | Genome | PX497241.1 |
| SARS-CoV-2 | BA.2.5 | Genome | PX105770.1 |
| SARS-CoV-2 | BA.1 | Amplicon | ERR7645572 |
| SARS-CoV-2 | BA.2.5 | Amplicon | ERR9236727 |
| SARS-CoV-2 | BA.2 | Amplicon | ERR9236855 |
| SARS-CoV-2 | B.1.617.2 | Amplicon | ERR13149562 |
| SARS-CoV-2 | AY.2 | Amplicon | SRR14854062 |
| SARS-CoV-2 | P.1 | Amplicon | ERR10012445 |
| SARS-CoV-2 | B.1.351 | Amplicon | ERR10159311 |
| SARS-CoV-2 | B.1.526 | Amplicon | ERR10741726 |
| Mosquito COX1 | *Culex australicus* | Genome | KU494997.1 |
| Mosquito COX1 | *Aedes notoscriptus* | Genome | NC_025473.1 |
| Mosquito COX1 | *Anopheles annulipes* | Genome | NC_054312.1 |
| Mosquito COX1 | *Coquillettidia linealis* | Genome | NC_054313.1 |
| Mosquito COX1 | *Coquillettidia xanthogaster* | Genome | OR509931.1 |
| Mosquito COX1 | *Culex annulirostris* | Genome | NC_054323.1 |
| Mosquito COX1 | *Culex australicus* | Genome | NC_054314.1 |
| Mosquito COX1 | *Culex sitiens* | Genome | NC_054318.1 |
| Mosquito COX1 | *Mansonia uniformis* | Genome | NC_047479.1 |
| Mosquito COX1 | *Mansonia uniformis* | Genome | OR001809.1 |
| Mosquito COX1 | *Aedes procax* | Genome | OQ946975.1 |
| Mosquito COX1 | *Aedes multiplex* | Genome | OR509933.1 |
| Mosquito COX1 | *Aedes notoscriptus* | Genome | PQ046711.1 |
| Mosquito COX1 | *Culex quinquefasciatus* | Genome | MN389462.1 |
| Mosquito COX1 | *Aedes procax* | Single species pool<br>COX1 capture | NA |
| Mosquito COX1 | *Aedes multiplex* | Single species pool<br>COX1 capture | NA |
| Mosquito COX1 | *Culex sitiens* | Single species pool<br>COX1 capture | NA |
| Mosquito COX1 | *Culex annulirostris* | Single species pool<br>COX1 capture | NA |
| *Mosquito COX1* | *Aedes notoscriptus* | Single species pool<br>COX1 capture | NA |
| *Mosquito COX1* | *Coquillettidia linealis* | Single species pool<br>COX1 capture | NA |
| *Mosquito COX1* | *Coquillettidia xanthogaster* | Single species pool<br>COX1 capture | NA |
| *Mosquito COX1* | *Anopheles annulipes* | Single species pool<br>COX1 capture | NA |
| *Mosquito COX1* | *Mansonia uniformis* | Single species pool<br>COX1 capture | NA |

